# N-Ethylmaleimide-Sensitive Factor Deletion in Dopamine D2 Receptor Cells and Associated Neuronal and Behavioural Changes in Mice

**DOI:** 10.64898/2025.12.17.694809

**Authors:** Min-Jue Xie, Koshi Murata, Hiroshi Kuniishi, Yugo Fukazawa, Noriyoshi Usui, Hideo Matsuzaki

## Abstract

N-ethylmaleimide-sensitive factor (NSF) plays a crucial role in neurotransmitter release and membrane molecule trafficking by regulating membrane fusion. Dysfunction of NSF has been linked to neuropsychiatric disorders. Although the interaction of NSF with dopamine receptor 2 (D2R) and the resulting excitotoxicity from its reduction *in vitro* have been demonstrated, the role of NSF in D2R-expressing cells *in vivo* remains unclear. This study investigated the effects of NSF loss on the survival of D2R-expressing cells and on mouse behaviour. We generated D2R-specific NSF conditional knockout (*Nsf*^f/f^;D2R-Cre) mice. Targeted deletion of NSF in D2R-expressing cells led to a significant decrease in D2R-expressing neurons, accompanied by increased apoptosis during postnatal development, reduced striatal volume, and substantially lowered dopamine levels in the striatum. Further evidence of dopaminergic dysfunction was shown by reduced dopamine transporter expression in the striatum and tyrosine hydroxylase expression in the striatum and substantia nigra. The *Nsf*^f/f^;D2R-Cre mice exhibited attention deficit hyperactivity disorder (ADHD)-like behaviours, including hyperactivity and impulsivity. Notably, combined administration of methylphenidate and a D2R agonist effectively alleviated hyperactivity and impulsivity, indicating a potential synergistic approach for ADHD treatment. These findings highlight the critical role of NSF in the survival of D2R-expressing neurons and suggest that disruption of the NSF-D2R interaction may contribute to ADHD-like phenotypes. This study underscores the translational relevance of the *Nsf*^f/f^;D2R-Cre model for ADHD and suggests that targeting D2R dysfunction, particularly in treatment-resistant cases, may represent a promising therapeutic strategy for ADHD.

## INTRODUCTION

N-ethylmaleimide-sensitive factor (NSF) is a homohexameric ATPase that plays an essential role in the protein machinery mediating membrane fusion events, including inter-cisternal Golgi protein transport and synaptic vesicle exocytosis [1–4]. Dysfunction of NSF has been associated with several neuropsychiatric disorders. Reduced NSF expression has been reported in lymphocytes of patients with autism spectrum disorder (ASD) and in the postmortem brains of patients with schizophrenia, whereas NSF-containing aggregates have been observed in basal ganglia specimens from individuals with Parkinson’s disease [5–7]. Moreover, genetic studies have linked SNARE complex genes, including those encoding NSF-interacting proteins, to attention deficit hyperactivity disorder (ADHD), ASD, major depressive disorder, bipolar disorder, and schizophrenia [8].

NSF interacts with various receptors, transporters, and neurotransmitter systems. Previously, we identified an interaction between NSF and the serotonin transporter (SERT), showing that NSF promotes SERT localisation to the cell membrane *in vitro* [5]. To validate this interaction *in vivo*, we generated *Nsf* heterozygous (*Nsf*^+/-^) mice, which showed reduced SERT and AMPA receptor expression at the membrane [9]. Additionally, these *Nsf*^+/-^ mice exhibited ASD-like behaviours, including impaired social communication, repetitive behaviours, and heightened anxiety.

NSF is abundantly expressed in the hippocampus and cerebral cortex, as well as in subcortical nuclei such as the striatum. *In vitro* studies have demonstrated that NSF interacts with dopamine receptors D1 (D1R) and D2 (D2R), regulating their membrane localisation [10, 11]. D2R activation has been shown to protect neurons against glutamate-induced excitotoxicity by initiating anti-apoptotic signalling via NSF [12, 13]. However, the neuroprotective role of the NSF-D2R interaction *in vivo* remains unclear.

D2R is expressed in several cell types, with particularly high levels in the medium spiny neurons of the striatum, which form a major component of the basal ganglia’s indirect pathway. This pathway plays a crucial role in motor control, inhibitory regulation, and reward processing [14]. Dysregulation of D2R function has been implicated in multiple neuropsychiatric disorders, including ADHD [15]. ADHD is characterised by difficulties in sustaining attention, excessive motor activity, and impulsivity, affecting approximately 1.4–3.0% of children and adolescents worldwide [16]. Neuroimaging studies have shown reduced availability of D2R/D3R in the striatum of individuals with ADHD, together with diminished dopamine transporter (DAT) expression and decreased striatal volume [15, 17]. These findings suggest that D2R dysfunction represents a central mechanism contributing to the neurobiological basis of ADHD.

Although D2R has been extensively studied, the upstream mechanisms regulating its function and localisation remain poorly understood. The interaction between NSF and D2R, particularly within striatal neurons, may play a crucial role in sustaining D2R-mediated signalling. To investigate this hypothesis, we generated D2R-specific *Nsf* conditional knockout (*Nsf*^f/f^;D2R-Cre) mice to examine the function of NSF in D2R-expressing cells *in vivo*.

## MATERIALS AND METHODS

### Animals

This study was reported in conformity with ARRIVE guidelines [18]. *Nsf*^f/f^;D2R-Cre mice and their control littermates (*Nsf*^f/f^) were used in this study. All experimental procedures were approved by the Animal Research Committee of University of Fukui and conducted in accordance with institutional guidelines and national regulations. Every effort was made to minimise the number of animals used and to reduce their suffering.

### Generation of NSF conditional knockout mice

To generate conditional *Nsf* knockout mice, a targeted *Nsf* allele was engineered by inserting cDNA into a lacZ-neo cassette [9]. This cassette was flanked by Frt sites, and the mice were subsequently crossed with B6;SJL-Tg (ACTFLPe) transgenic mice (The Jackson Laboratory, Bar Harbor, ME, USA) to excise the neomycin resistance gene. This procedure produced floxed *Nsf* (*Nsf^f/+^*) mice.

Homozygous *Nsf*^f/f^ mice were then crossed with *D2R-Cre* mice to generate independent lines of *Nsf*^f/f^;D2R-Cre mice. The *D2R-Cre* mouse line used in this study, B6.FVB(Cg)-Tg(Drd2-Cre)ER44Gsat/Mmucd (RRID: MMRRC_032108-UCD), was obtained from the Mutant Mouse Resource and Research Centre (MMRRC) at the University of California, Davis, an NIH-funded repository. This strain was originally developed by Nathaniel Heintz, Ph.D. (The Rockefeller University, GENSAT) and Charles Gerfen, Ph.D. (National Institute of Mental Health, NIH) and donated to the MMRRC [19].

### *In situ* hybridisation

To detect *Nsf*, *D1R*, or *D2R* expression, *in situ* hybridisation was performed using DIG-labelled antisense RNA probes. A plasmid for Nsf probe synthesis was constructed using the pGEM-T kit (Promega, Tokyo, Japan) and an amplified PCR product with the following primers: CTTGTCTTTAGCTTCAATGATAA-CGATAAGATTGAGCGACGAA (183 bp). Plasmid templates for Drd1 and Drd2 probes were kindly provided by Dr Kazuto Kobayashi [20]. The experimental procedures followed previously established methods [21]. ImageJ software (NIH, Bethesda, MD, USA) was used to analyse and quantify stained cells.

### Immunohistochemistry (IHC) and western blot (WB)

IHC and WB experiments were performed as described by Xie et al. [22]. The following primary antibodies were used: anti-D2R (AB_2571596, Frontier Institute Co., Hokkaido, Japan), anti-DAT (AB_2571688, Frontier Institute Co.), anti-TH (MAB318, Merck Ltd., Darmstadt, Germany), anti-NeuN (ab177487, Abcam, Cambridge, UK), anti-NeuN (MAB377, Merck Ltd.), anti-single-stranded DNA (ssDNA) (18731, Immuno-Biological Laboratories Co., Gunma, Japan), and anti-HRP-GAPDH antibody (M171-7, MBL, Toyama, Japan). Quantification of NeuN- and TH-immunopositive cells was performed using ImageJ software.

### Quantitative real-time reverse-transcription-polymerase chain reaction (qRT-PCR)

Total RNA was extracted from the striatum and reverse-transcribed into cDNA using the GeneAce cDNA Synthesis Kit (Nippon Gene co., Tokyo, Japan) following the manufacturer’s instructions. qRT-PCR was performed using SYBR Green Master Mix (Thermo Fisher Scientific, Waltham, MA, USA) on a QuantStudio 5 Real-Time PCR System (Thermo Fisher Scientific). The following primer pairs were used for target gene amplification: D2R, 5′-GCAGCCGAGCTTTCAGAGCC and 5′**-**GGGATGTTGCAGTCACAGTG; Cre, 5′**-**CTGATTTCGACCAGGTTCGT and 5′-ATTCTCCCACCGTCAGTACG; GAPDH, 5′-ACTCCACTCACGGCAAATTC and 5′**-**TCTCCATGGTGGTGAAGACA [23].

### Stereotaxic surgery and adeno-associated virus (AAV) injection in the striatum

Stereotaxic surgery and AAV injection were performed as previously described [24]. Mice were anaesthetised by intraperitoneal injection of a mixture containing medetomidine (0.75 mg/kg), midazolam (4 mg/kg), and butorphanol (5 mg/kg), and secured in a stereotaxic frame (Narishige, Tokyo, Japan). A mixture of viral vectors, AAV5- human synapsin (hSyn)-EGFP (#50465, titre: 8.4 × 10^12^ vg/mL) and AAV5-hSyn-DIO-mCherry (#50459, titre: 8.4 × 10^12^ vg/mL) (Addgene, Watertown, USA) was combined at a 1:1 ratio and injected into the striatum (depth: 2.0 mm from the dura; volume: 300 nL; infusion rate: 100 nL/min) using a 10 μL Hamilton syringe connected to an infusion pump (UMP-3, World Precision Instruments, FL, USA). More information is provided in the Supplementary Methods.

### Measurement of dopamine concentrations

Dopamine concentrations were quantified by Eicom Co., Ltd. using high-performance liquid chromatography (HPLC) equipped with an electrochemical detector (ECD-100, Eicom, Kyoto, Japan). Striatal tissues were dissected from mice, rapidly frozen in liquid nitrogen, and stored at -80 °C until analysis. The subsequent procedures followed previously established methods [25].

### Behavioural tests

Behavioural tests were performed during the light phase of the illumination cycle (between 10:00 a.m. and 4:00 p.m.). Each mouse was subjected to only one behavioural test per day. Details of the behavioural battery, including the open-field test [9], cliff avoidance and jumping test [25, 26], elevated plus maze [24], prepulse inhibition (PPI) test, and three-chamber social interaction test [9], are provided in the Supplementary Methods.

### Statistical analysis

All statistical analyses were conducted using IBM SPSS Statistics version 23 and JMP Pro version 14. Data were analysed using either a two-tailed Student’s *t*-test. For the analysis of average striatal area, a two-way repeated-measures analysis of variance (ANOVA) with Tukey’s post-hoc test was applied. Statistical significance was set at *p*<0.05 for all tests.

## RESULTS

### Generation of *Nsf*^f/f^;D2R-Cre mice

To overcome the embryonic lethality of conventional *Nsf*-null mice and investigate the role of NSF in D2R-expressing cells, we generated a conditional D2R-specific *Nsf* knockout line. Using a targeting vector from the KOMP Repository (Vector ID: PG00174_Z_5_D06), a floxed allele was inserted between exons 5 and 6 of the *Nsf* gene (Fig. 1a) [9]. Mice carrying the targeted allele, flanked by *Frt* sequences, were first bred with transgenic mice expressing Flp recombinase to generate *Nsf*^f/f^ mice [27]. These *Nsf*^f/f^ mice were then crossed with D2R-Cre BAC transgenic mice [19] to create *Nsf*^f/f^;D2R-Cre mice, which exhibit NSF disruption specifically in D2R-expressing cells. Successful targeting of the allele was confirmed via genotyping using PCR analysis (Fig. 1b).

**Fig. 1:**
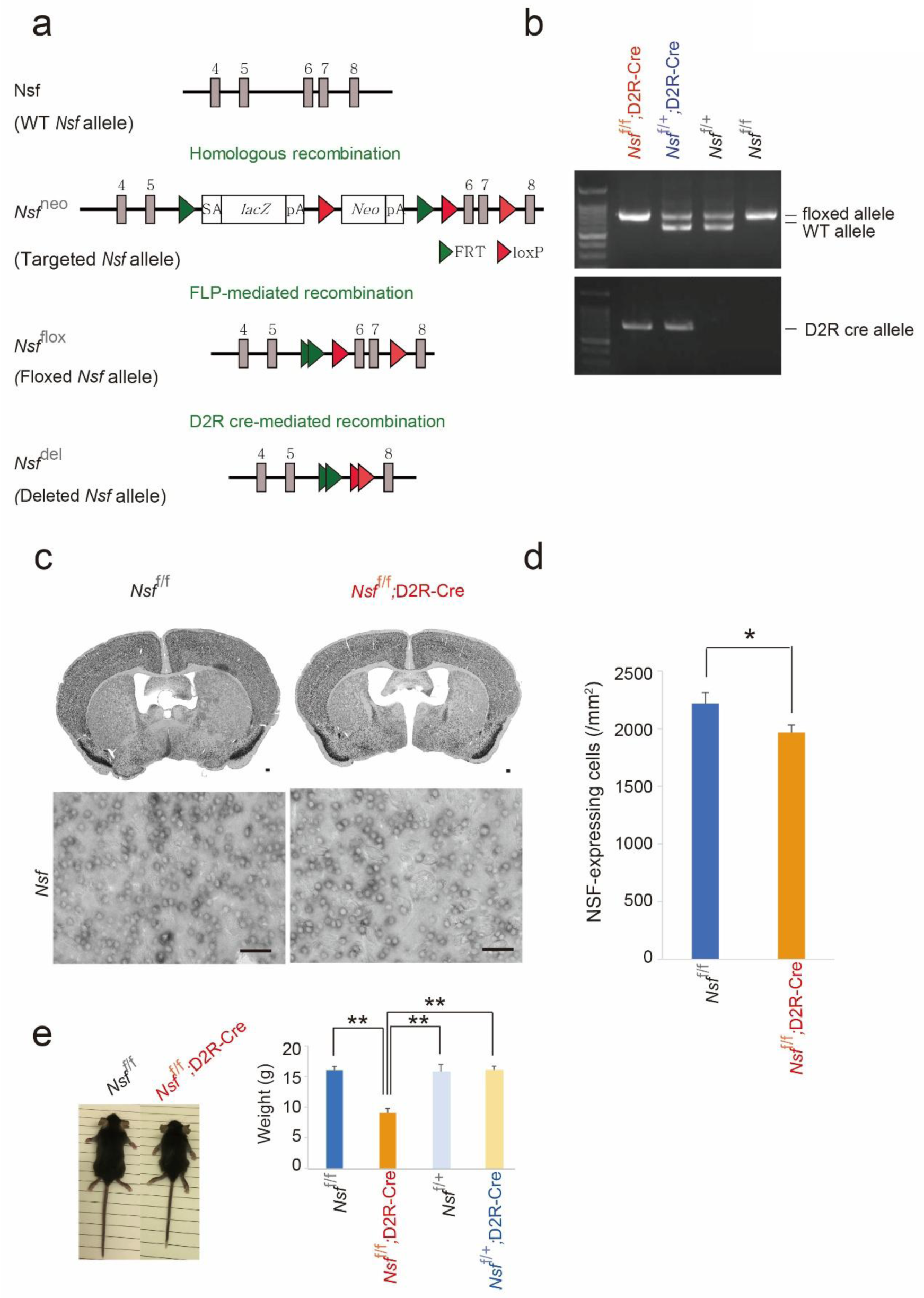
Conditional knockout of *Nsf* in D2R-expressing cells in mice. (a) Schematic representation of the knockout-first conditional allele. The targeting strategy involves the removal of the neomycin resistance cassette at the *Nsf* locus. Exons 6 and 7 of the *Nsf* gene are floxed, enabling their removal upon Cre recombinase induction (SA, splice acceptor; Laz, β-gal and neomycin phosphotransferase fusion gene; pA, polyadenylation signal sequence; Neo, neomycin resistance cassette; FRT, Frt sites; Lox, loxP sites). Floxed *Nsf* mice are crossed with D2R-Cre mice. This cross allows for the conditional deletion of the *Nsf* gene specifically in D2R-expressing cells. (b) PCR detection of floxed, wild-type (WT), and D2R-Cre alleles. (c) *In situ* hybridisation showing *Nsf* mRNA expression in the striatum. The upper row displays a scale bar representing 500 µm, while the lower row presents a high-magnification image with a scale bar representing 50 µm. (d) The number of *Nsf* mRNA-expressing cells per mm² in the striatum was quantified in *Nsf*^f/f^ and *Nsf*^f/f^;D2R-Cre mice at 3 weeks of age (n=6 for each genotype). (e) Representative mouse images and the weight of male mice at 3 weeks of age (*Nsf*^f/f^ mice: n=18; *Nsf*^f/f^;D2R-Cre mice: n=15; *Nsf*^f/+^;D2R-Cre mice: n=7; *Nsf*^f/+^ mice: n=7). Data are presented as mean±SEM. Student’s *t*-test; \*\**p*<0.01 (e), \**p*<0.05 (d).

To verify the disruption of *Nsf* expression in D2R-expressing striatal cells, we performed *in situ* hybridisation using an antisense probe targeting *Nsf* mRNA. The number of *Nsf*-expressing cells in the striatum was significantly reduced in *Nsf*^f/f^;D2R-Cre mice compared with that in *Nsf*^f/f^ controls (*Nsf*^f/f^ mice: 2 220.1±93.8 cells/mm^2^; *Nsf*^f/f^;D2R-Cre mice: 1 965.3±65.9 cells/mm^2^; Student’s *t*-test, *p*=0.047; Fig. 1c,d). Furthermore, *Nsf*^f/f^;D2R-Cre mice exhibited significant postnatal weight loss compared with that in all control groups. At postnatal day 21 (P21), body weight was markedly lower in *Nsf*^f/f^;D2R-Cre mice (9.0±0.7 g) than in controls (*Nsf*^f/f^ mice: 16.0±0.6 g; *Nsf*^f/+^;D2R-Cre mice: 16.0±0.7 g; *Nsf*^f/+^ mice: 15.7±1.3 g; Student’s *t*-test, *p*<0.01; Fig. 1e).

### Decrease in the number of striatal D2R-expressing cells in *Nsf*^f/f^;D2R-Cre mice

We next conducted qRT-PCR analysis to measure *D2R* mRNA expression in the striatum. A significant reduction was observed in *Nsf*^f/+^;D2R-Cre mice (*Nsf*^f/f^ mice: 0.69±0.10; *Nsf*^f/+^;D2R-Cre mice: 0.75±0.07; *Nsf*^f/f^;D2R-Cre mice: 0.02±0.002; Student’s *t*-test, *p*<0.001; Fig. 2a). We then examined the number of *D2R*-expressing cells in *Nsf*^f/f^;D2R-Cre mice using *in situ* hybridisation, including *D1R*-expressing cells as a Cre-negative control population. The number of *D2R*-expressing cells per unit area was significantly reduced in *Nsf*^f/f^;D2R-Cre mice compared with that in *Nsf*^f/f^ control (*Nsf*^f/f^ mice: 1 477.8±59.2 cells/mm^2^; *Nsf*^f/f^;D2R-Cre mice: 102.2±54.1 cells/mm^2^; Student’s *t*-test, *p*<0.001; Fig. 2b,d). Similarly, the proportion of *D2R*-expressing cells relative to total cells was markedly decreased in *Nsf*^f/f^;D2R-Cre mice (*Nsf*^f/f^ mice: 60.0%±2.69%; *Nsf*^f/f^;D2R-Cre mice: 3.32%±2.24%; Student’s *t*-test, *p*<0.001; Fig. 2e). In contrast, the number of D1R-expressing cells per unit area was significantly increased in *Nsf*^f/f^;D2R-Cre mice (*Nsf*^f/f^ mice: 1 333.3±74.6 cells/mm^2^; *Nsf*^f/f^; D2R-Cre mice: 2 145.9±139.9 cells/mm^2^; Student’s *t*-test, *p*<0.001; Fig. 2c,f). Similarly, the proportion of D1R-expressing cells relative to total cells was elevated in *Nsf*^f/f^;D2R-Cre mice (*Nsf*^f/f^ mice: 65.4±2.69%; *Nsf*^f/f^; D2R-Cre mice: 93.42±1.27%; Student’s *t*-test, *p*<0.001) (Fig. 2g). Striatal cell density did not differ significantly between the two groups (*Nsf*^f/f^ mice: 2 304.4±81.4 cells/mm^2^, *Nsf*^f/f^;D2R-Cre mice: 2 234.0±85.6 cells/mm^2^; Student’s *t*-test, *p*=0.57; Fig. 2h), suggesting that overall striatal shrinkage may be attributable to the selective loss of D2R-expressing cells.

**Fig. 2:**
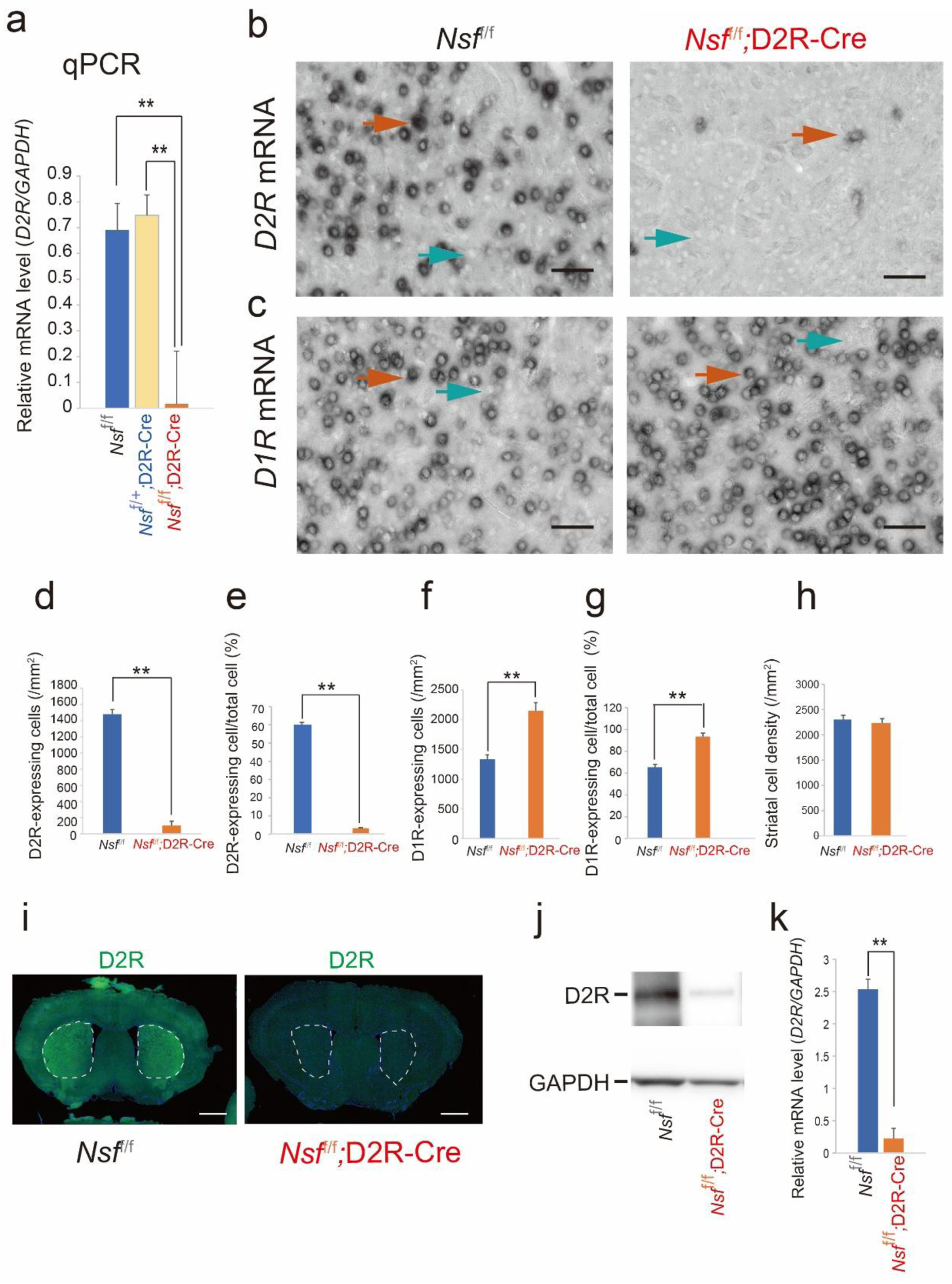
Reduced striatal *D2R* expression in *Nsf*^f/f^;D2R-Cre mice. (a) The *D2R* mRNA expression level was analysed using qPCR, normalised to *Gapdh*, in the striatum of *Nsf*^f/f^ and *Nsf*^f/f^;D2R-Cre mice at 3 months (n=7 for each genotype). (b) *D2R* and (c) *D1R* mRNA expression levels were analysed using *in situ* hybridisation in *Nsf*^f/f^ and *Nsf*^f/f^;D2R-Cre mice. Red arrows indicate cells expressing the gene, while green arrows indicate cells not expressing the gene. Scale bar=50 µm. The number of D2R-expressing cells per mm² (d) and per total cells (e) in the striatum was quantified in *Nsf*^f/f^ and *Nsf*^f/f^;D2R-Cre mice at 3 weeks of age (n=11 for each genotype). The number of D1R- expressing cells per mm² (f) and per total cells (g) in the striatum was quantified in *Nsf*^f/f^ and *Nsf*^f/f^;D2R-Cre mice at 3 weeks of age (n=11 for each genotype). (h) Quantification of the total cell count per mm^2^ in the striatum of *Nsf*^f/f^ (n=29) and *Nsf*^f/f^;D2R-Cre (n=27) mice. (i) Immunohistochemical staining for D2R in the striatum of *Nsf*^f/f^ (left panel) and *Nsf*^f/f^;D2R-Cre mice (right panel) at 3 weeks of age. Scale bars=1 mm. Results are representative of more than three independent experiments. (j) The D2R protein expression level was analysed via western blot, normalised to *Gapdh*, in the striatum of *Nsf*^f/f^ and *Nsf*^f/f^;D2R-Cre mice. (k) Quantification of D2R protein levels, normalised to GAPDH as internal reference, in the striatum of *Nsf*^f/f^ (n=7) and *Nsf*^f/f^;D2R-Cre (n=6) mice at postnatal day 8. Data are presented as mean±SEM. Student’s *t*-test; \*\**p*<0.01 (a, d-g, k).

We further confirmed the reduction in D2R expression at the protein level using immunofluorescence staining (Fig. 2i) and WB analysis (Fig. 2j). D2R protein levels were significantly lower in *Nsf*^f/f^;D2R-Cre mice (*Nsf*^f/f^ mice: 2.54±0.28; *Nsf*^f/f^;D2R-Cre mice: 0.22±0.08; Student’s *t*-test, *p*<0.001; Fig. 2j,k). These findings collectively demonstrate a significant reduction in both D2R mRNA and protein expression in *Nsf*^f/f^;D2R-Cre mice.

### Loss of Cre-expressing cells in the striatum of *Nsf*^f/f^;D2R-Cre mice

We then investigated two possible explanations for the reduction in both the number of D2R-expressing cells and D2R expression in the striatum: (1) downregulation of D2R expression within individual cells, or (2) actual loss of D2R-expressing cells. To distinguish between these possibilities, we assessed *Cre* mRNA expression. If D2R-expressing cells remained, they would continue to express *Cre.* We observed minimal *Cre* expression in *Nsf*^f/f^;D2R-Cre mice, whereas strong *Cre* expression was detected in *Nsf*^f/+^;D2R-Cre mice (*Nsf*^f/f^ mice: 0±0; *Nsf*^f/+^;D2R-Cre mice: 0.59±0.74; *Nsf*^f/f^;D2R-Cre mice: 0.01±0.005; Student’s *t*-test, *p*<0.001; Supplementary Fig. 1a,b). These findings suggest that the reduction in D2R expression results from the loss of D2R-expressing cells.

To further confirm the loss of *Cre*-expressing cells in *Nsf*^f/f^;D2R-Cre mice, we injected a mixture of two AAV vectors into the striatum: one carrying EGFP under the hSyn promoter and the other carrying *Cre*-dependent mCherry under the same promoter. This design enabled labelling of *Cre*-expressing cells with mCherry and all infected cells with EGFP. Following AAV injection, approximately 50–60% of EGFP-labelled cells expressed mCherry, indicating putative D2R-expressing cells, in control mice (*Nsf*^+/+^;D2R-Cre mice: 56±6%; *Nsf*^f/+^;D2R-Cre mice: 69±4%). In contrast, only a small proportion of EGFP-labelled cells expressed mCherry in *Nsf*^f/f^;D2R-Cre (6±2%; Student’s *t*-test, *p*<0.001; Fig. 3a,b.c). These results indicate that the decreased D2R mRNA and protein expression observed in *Nsf*^f/f^;D2R-Cre mice stems from the loss of D2R-expressing cells.

**Fig. 3:**
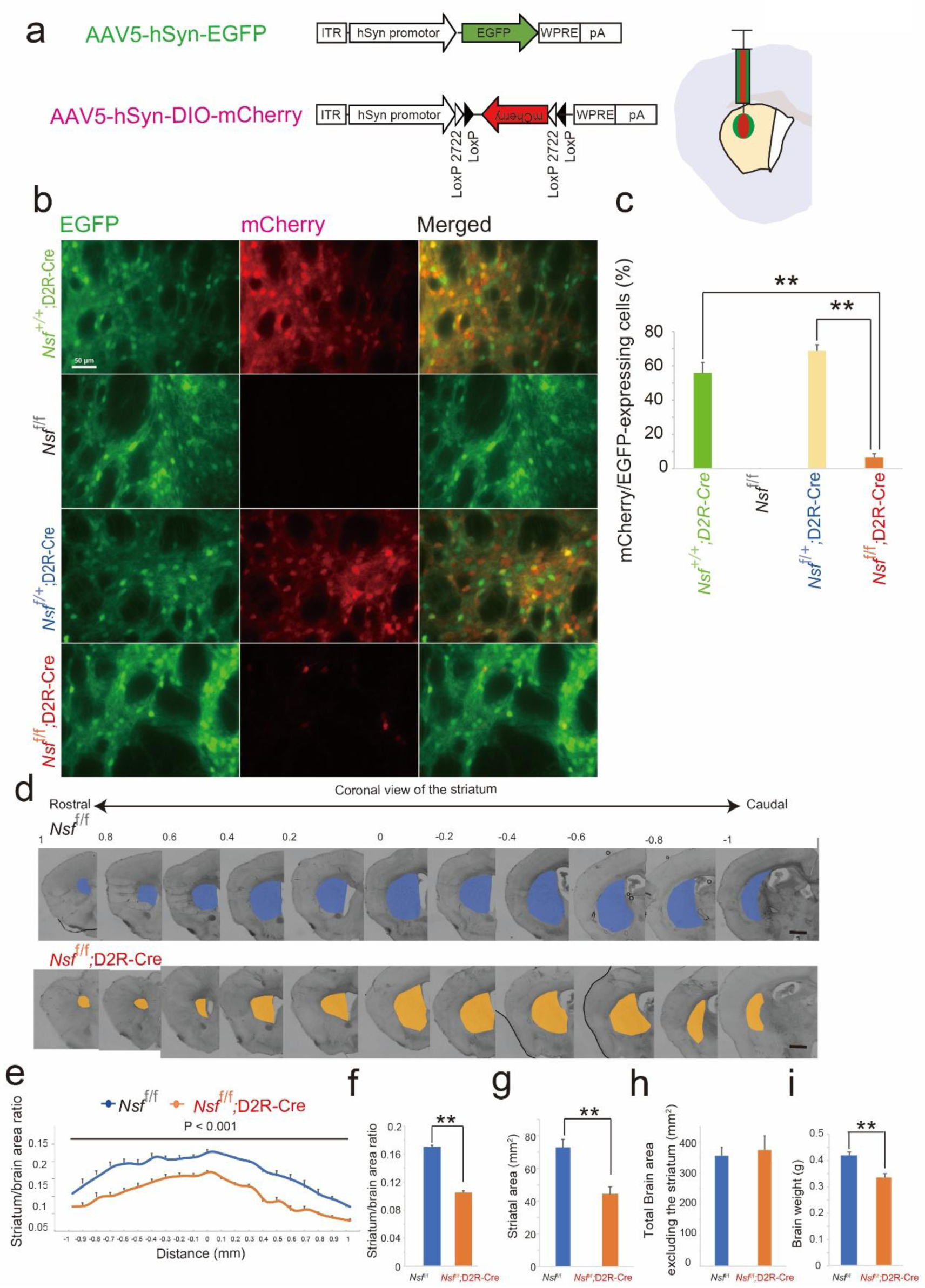
Loss of striatal D2R-expressing cells and shrinkage of the striatum in *Nsf*^f/f^;D2R-Cre mice. (a) Schematic illustration of the adeno-associated virus (AAV) constructs: AAV5-hSyn-EGFP, in which the human synapsin promoter (hSyn) drives EGFP expression, and AAV5-hSyn-DIO-mCherry, a Cre-dependent construct expressing mCherry. AAVs were injected into the striatum at 4 weeks of age and analysed at 5 weeks. (b) Representative images of the striatum showing EGFP and mCherry fluorescence in *Nsf*^+/+^;D2R-Cre, *Nsf*^f/f^, *Nsf*^f/+^;D2R-Cre, and *Nsf*^f/f^;D2R-Cre mice. Scale bars=50 μm. (c) Quantification of mCherry-expressing cells relative to EGFP-expressing cells in the striatum (*Nsf*^f/f^ mice: n=3; *Nsf*^f/f^;D2R-Cre mice: n=4; *Nsf*^f/+^;D2R-Cre mice: n=5; *Nsf*^f/+^ mice: n=7). (d) Representative coronal brain sections through the anterior commissure (0 axial level) from 3-week-old *Nsf*^f/f^ (upper panel) and *Nsf*^f/f^;D2R-Cre (lower panel) mice. (e) Average striatal area normalised to total brain area in *Nsf*^f/f^ and *Nsf*^f/f^;D2R-Cre mice (n=8 per group). Scale bars=500 μm. (f) Total striatal area relative to total brain area in *Nsf*^f/f^ and *Nsf*^f/f^;D2R-Cre mice (n=6 per genotype). (g) Absolute striatal area (n=6 per genotype). (h) Total brain area excluding the striatum in *Nsf*^f/f^ and *Nsf*^f/f^;D2R-Cre mice (n=6 per genotype). (i) Brain weight of *Nsf*^f/f^ and *Nsf*^f/f^;D2R-Cre mice (n=8 per genotype). Data are presented as mean±SEM. Student’s *t*-test (c, f-i) and two-way repeated measures ANOVA, F(1,20)=2.663, *p*=0.001 (e); \*\**p*<0.01 (c, f, g, i).

### Increased cell death and shrinkage of the striatum in *Nsf*^f/f^;D2R-Cre mice

We next investigated whether cell death during postnatal development contributes to the loss of striatal D2R-expressing cells in *Nsf*^f/f^;D2R-Cre mice. To assess this, we used ssDNA as a sensitive and specific marker of cell death. At postnatal day 8 (P8), the number of ssDNA-positive cells in the striatum was significantly higher in *Nsf*^f/f^;D2R-Cre mice than in *Nsf*^f/f^ controls (*Nsf*^f/f^ mice: 5.68 ±1.19 cells/mm^2^; *Nsf*^f/f^;D2R-Cre mice: 13.28±1.70 cells/mm^2^; Student’s *t*-test, *p*=0.013; Supplementary Fig. 2a,c). However, by P21, almost no ssDNA-positive cells were observed in either group (*Nsf*^f/f^ mice: 0.46±0.12 cells/mm^2^; *Nsf*^f/f^;D2R-Cre mice: 0.24±0.10 cells/mm^2^; Student’s *t*-test, *p*=0.35; Supplementary Fig. 2b,c), suggesting that NSF exerts a protective effect on D2R-expressing cells during postnatal development.

We then evaluated whether the striatal volume was reduced in *Nsf*^f/f^;D2R-Cre mice as a consequence of cell death. The ratio of striatal area to whole brain area was measured using coronal sections encompassing the entire striatum. This ratio was significantly reduced in *Nsf*^f/f^;D2R-Cre mice compared with that in *Nsf*^f/f^ controls (two-way repeated measures ANOVA, F (1,20)=2.663, *p*=0.001; Fig. 3d.e). Specifically, the striatal-to-whole-brain ratio decreased from 0.17 in *Nsf*^f/f^ mice to 0.11 in *Nsf*^f/f^;D2R-Cre mice (*Nsf*^f/f^ mice: 0.17±0.002; *Nsf*^f/f^;D2R-Cre mice: 0.11±0.002; Student’s *t*-test, *p*<0.001; Fig. 3f). The total striatal area was also significantly smaller in *Nsf*^f/f^;D2R-Cre mice (*Nsf*^f/f^ mice: 72.9±4.8 mm^2^; *Nsf*^f/f^;D2R-Cre mice: 44.1±4.6 mm^2^; Student’s *t-*test, *p*=0.005; Fig. 3g). In contrast, the total brain area excluding the striatum did not differ significantly between the two groups (*Nsf*^f/f^ mice: 355.8±27.8 mm^2^; *Nsf*^f/f^;D2R-Cre mice: 374.4±45.6 mm^2^; Student’s *t*-test, *p*=0.75; Fig. 3h). Additionally, brain weight of *Nsf*^f/f^;D2R-Cre mice was significantly reduced compared with that of *Nsf*^f/f^ controls (*Nsf*^f/f^ mice: 0.42±0.01; *Nsf*^f/f^;D2R-Cre mice: 0.34±0.01; Student’s *t*-test, *p*<0.001; Fig. 3i). These findings demonstrate that increased striatal cell death is accompanied by regional shrinkage, implicating NSF as a key factor in maintaining striatal integrity during postnatal development.

### Decreased striatal dopamine levels in *Nsf*^f/f^;D2R-Cre mice

We examined whether striatal dopamine levels were affected in *Nsf*^f/f^;D2R-Cre mice using HPLC-ECD. Dopamine concentration in the striatum was reduced by approximately 90% in *Nsf*^f/f^;D2R-Cre mice compared with that in *Nsf*^f/f^ mice at P21 (*Nsf*^f/f^;D2R-Cre mice: 0.30±0.02 ng/mg tissue; *Nsf*^f/f^ mice: 3.97±0.040 ng/mg tissue; Student’s *t-*test, *p*<0.001; Fig. 4a).

**Fig. 4:**
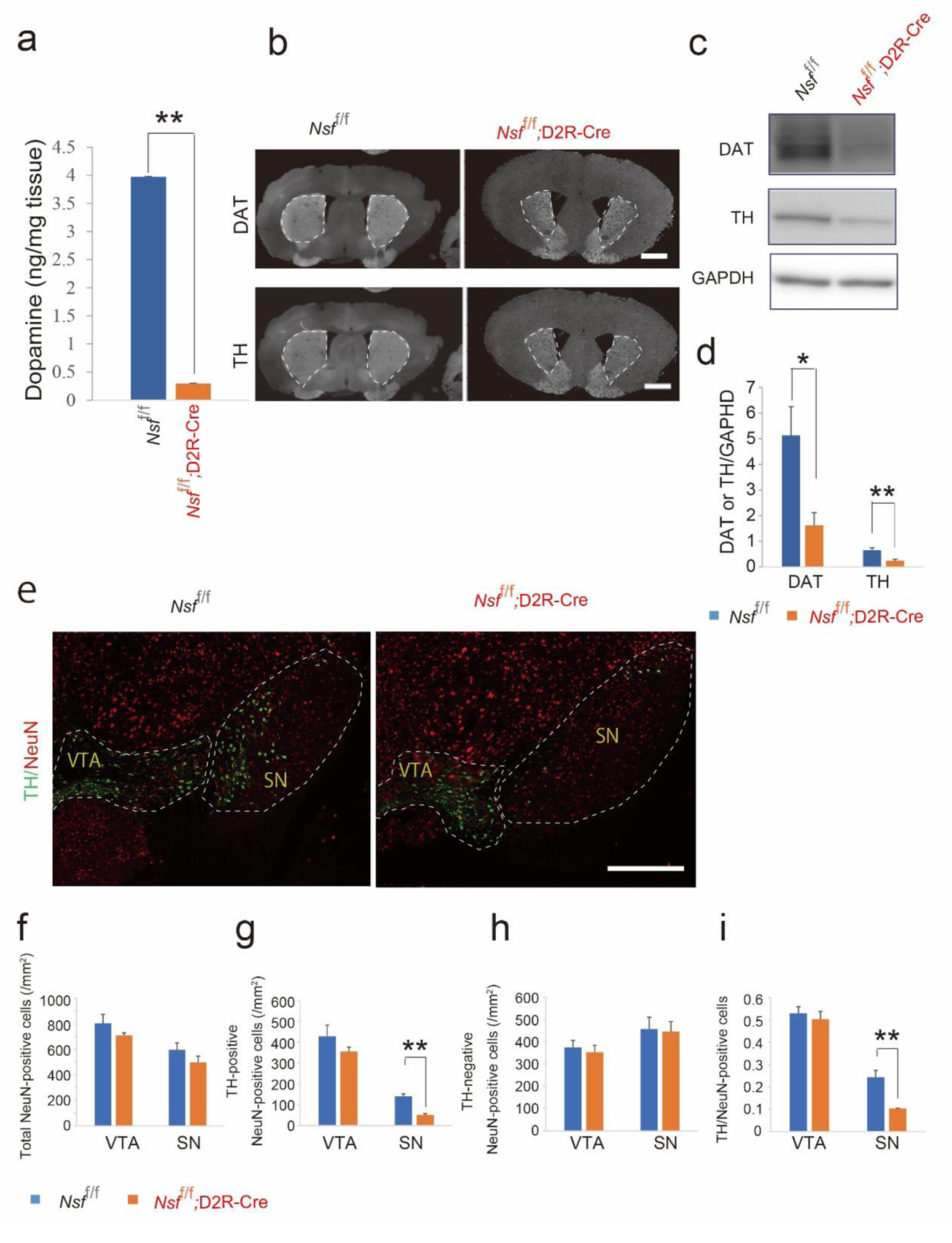
Decreased striatal dopamine level and DAT and TH expression in *Nsf*^f/f^;D2R-Cre mice. (a) Dopamine, serotonin, and norepinephrine levels in the striatal tissue from *Nsf*^f/f^ and *Nsf*^f/f^;D2R-Cre mice at P21, measured via HPLC-ECD (n=5 per genotype). Data are presented as mean±SEM. Student’s *t-*test, \*\**p*<0.01. (b) Immunofluorescence detection of DAT and TH expression in the striatum of 4-week-old *Nsf*^f/f^ and *Nsf*^f/f^;D2R-Cre mice. Scale bars=500 μm. (c) Western blot analysis of DAT, TH, and GAPDH (internal control) expression levels. (d) Quantification of relative band densities for DAT and TH in the striatum of *Nsf*^f/f^ and *Nsf*^f/f^;D2R-Cre mice via scanning densitometry (n=7 per genotype). (e) Immunofluorescence detection of TH and NeuN expression in the substantia nigra (SN) and ventral tegmental area (VTA) of 4-week-old *Nsf*^f/f^ and *Nsf*^f/f^;D2R-Cre mice. Scale bars=1 mm. (f-i) Quantification of NeuN-positive cells (f), TH-positive cells (g), TH-negative cells (h), and the ratio of TH-positive to NeuN-positive cells (TH/NeuN) (i) in the SN and VTA (n=4 per genotype). Data are presented as mean±SEM. Student’s *t*-test; \*\**p*<0.01 (g, i).

We next assessed the expression of DAT and TH, which regulate dopamine neurotransmission involved in reuptake and biosynthesis, respectively [29]. Immunofluorescence and WB analyses revealed markedly reduced DAT and TH expression in the striatum of *Nsf*^f/f^;D2R-Cre mice compared with that of *Nsf*^f/f^ controls (for DAT, *Nsf*^f/f^ mice: 5.13±1.11; *Nsf*^f/f^;D2R-Cre mice: 1.63±0.49; Student’s *t*-test, *p*=0.019; for TH, *Nsf*^f/f^ mice: 0.66±0.09; *Nsf*^f/f^;D2R-Cre mice: 0.24±0.0.05; Student’s *t-*test, *p*=0.003; Fig. 4b-d).

We further examined whether TH expression was similarly reduced in the midbrain, where dopaminergic neurons originate and project their axons to the striatum. TH expression was significantly decreased in the substantia nigra (SN) of *Nsf*^f/f^;D2R-Cre mice compared with that in *Nsf*^f/f^ controls (TH-positive cells: *Nsf*^f/f^ mice: 140.1±11.8 mm^2^; *Nsf*^f/f^;D2R-Cre mice: 51.2±6.1 mm^2^; Student’s *t*-test, *p*=0.001; Fig. 4e,g). In contrast, the number of TH-negative cells in the SN did not differ significantly between the groups (*Nsf*^f/f^ mice: 456.2±52.6 mm^2^; *Nsf*^f/f^;D2R-Cre mice: 445.0±45.3 mm^2^; *p*=0.89; Fig. 4 e,h). No significant differences were observed in the ventral tegmental area (VTA) for either TH-positive cells (*Nsf*^f/f^ mice: 428.3±53.4 mm^2^; *Nsf*^f/f^;D2R-Cre mice: 355.3±21.3 mm^2^; *p*=0.31) or TH-negative cells (*Nsf*^f/f^ mice: 376.6±31.4 mm^2^; *Nsf*^f/f^;D2R-Cre mice: 353.0±30.4 mm^2^; *p*=0.68) (Fig. 4e,g,h). Total neuronal counts, assessed via NeuN immunolabelling, showed no significant differences between the groups in either the SN or VTA (SN: *Nsf*^f/f^ mice, 596.3±53.4 mm^2^; *Nsf*^f/f^;D2R-Cre mice, 496.2±51.3 mm^2^; *p*=0.29. VTA: *Nsf*^f/f^ mice, 802.9±70.7 mm^2^; *Nsf*^f/f^;D2R-Cre mice: 708.3±21.2 mm^2^; *p*=0.31) (Fig. 4e,f). The ratio of TH-positive to NeuN-positive cells (TH/NeuN) was significantly lower in the SN of *Nsf*^f/f^;D2R-Cre mice (*Nsf*^f/f^ mice: 0.24±0.03; *Nsf*^f/f^;D2R-Cre mice: 0.10±0.002; *p*=0.007), while no difference was observed in the VTA (*Nsf*^f/f^ mice: 0.53±0.03; *Nsf*^f/f^;D2R-Cre mice: 0.50±0.03; *p*=0.64) (Fig. 4e,i). These findings suggest that *Nsf*^f/f^;D2R-Cre mice exhibit markedly reduced striatal dopamine levels, accompanied by decreased TH expression in dopaminergic neurons of the SN.

### Hyperactivity and abnormal impulsive behaviour in *Nsf*^f/f^;D2R-Cre mice

To investigate the behavioural characteristics of *Nsf*^f/f^;D2R-Cre mice, we conducted a comprehensive behavioural test battery in mice over 6 months of age. These mice exhibited pronounced hyperactivity, as indicated by a significant increase in total locomotor distance compared with that in *Nsf*^f/f^ controls (*Nsf*^f/f^ mice: 16 934±1 531 cm; *Nsf*^f/f^;D2R-Cre mice: 75 831±8 808cm; Student’s t-test, *p*<0.001; Fig. 5a). In addition to hyperactivity, *Nsf*^f/f^;D2R-Cre mice displayed impulsive-like behaviours, a hallmark of ADHD. Impulsivity was assessed using the cliff-avoidance test, which evaluates a rodent’s tendency to jump from an elevated platform as an indicator of impulsive-like behaviour [26, 27]. Impulsivity was elevated in *Nsf*^f/f^;D2R-Cre mice, as reflected by significantly increased jumping activity (*Nsf*^f/f^ mice: 362.9±33.6 s; *Nsf*^f/f^;D2R-Cre mice: 77.4±40.5 s; Student’s t-test, *p*<0.001; Fig.5c). During the test, 60% *Nsf*^f/f^;D2R-Cre mice and 0% *Nsf*^f/f^ mice jumped off the platform within the first minute. By the end of the 7-min session, 90% of *Nsf*^f/f^;D2R-Cre mice and only 25% of *Nsf*^f/f^ mice had jumped (Fig. 5d). Repetitive behaviour, assessed via vertical activity, did not differ significantly between the groups (*Nsf*^f/f^ mice: 367±43 time; *Nsf*^f/f^;D2R-Cre mice: 527±90 time; Student’s *t-*test, *p*=0.12; Fig. 5b). In the elevated plus maze test, *Nsf*^f/f^;D2R-Cre mice spent significantly more time in the closed arms than did *Nsf*^f/f^ mice (*Nsf*^f/f^ mice: 404±43 s; *Nsf*^f/f^;D2R-Cre mice: 507±15 s; Student’s *t*-test, *p*=0.046).

**Fig. 5:**
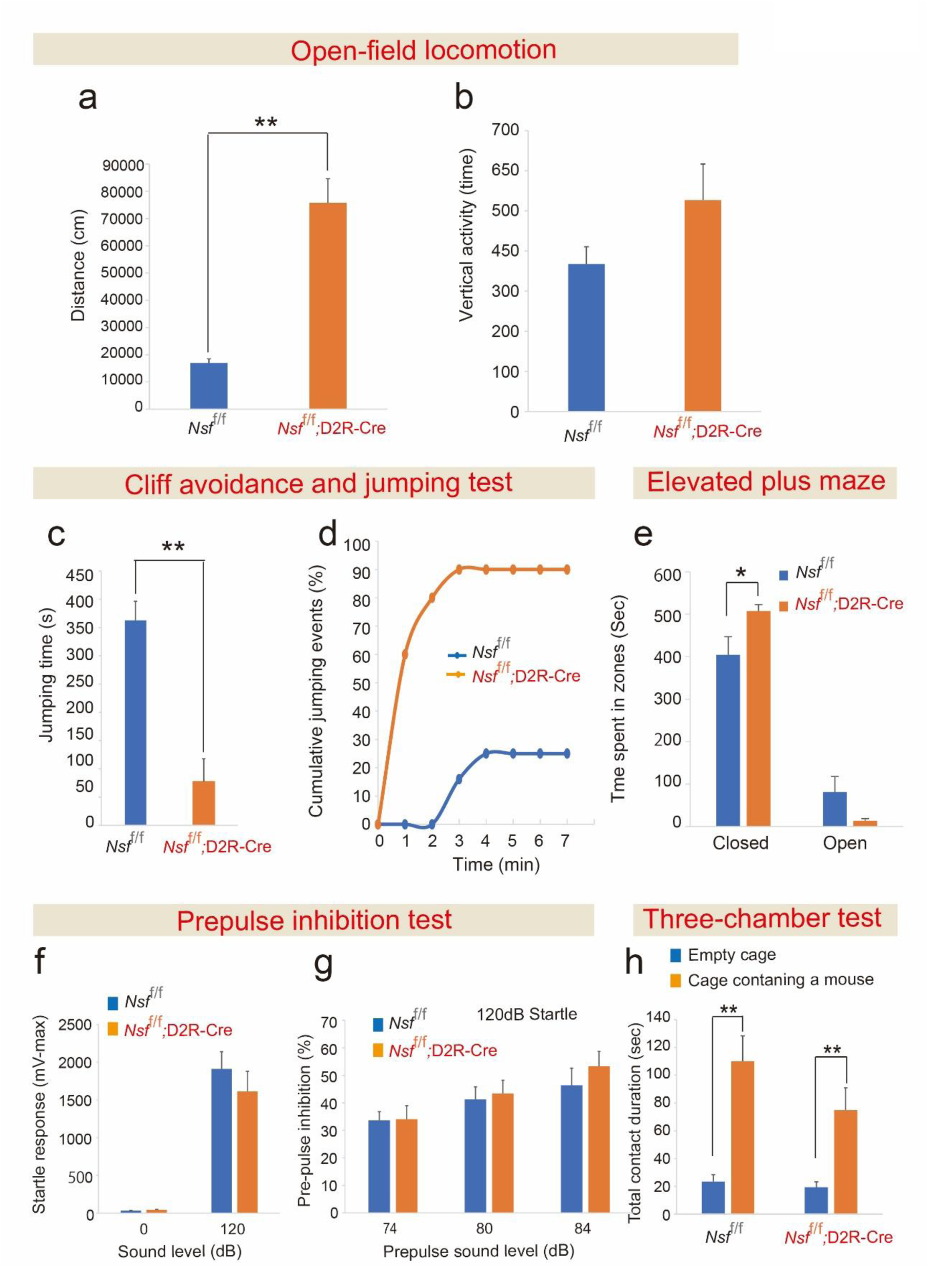
Hyperactivity and impulsivity in adult *Nsf*^f/f^;D2R-Cre mice. (a) Total distance travelled in the open-field test by adult *Nsf*^f/f^;D2R-Cre mice over 6 months of age (*Nsf*^f/f^: n=13; *Nsf*^f/f^;D2R-Cre: n=11). (b) Vertical activity in the open field, indicating repetitive behaviour (*Nsf*^f/f^: n=13; *Nsf*^f/f^;D2R-Cre: n=11). (c) Latency to the first fall in the cliff-avoidance reaction test (*Nsf*^f/f^: n=12; *Nsf*^f/f^;D2R-Cre: n=10). (d) Cumulative frequency of cliff-jumping events over 7 min (*Nsf*^f/f^: n=12; *Nsf*^f/f^;D2R-Cre: n=10). (e) Time spent in the open and closed arms during the elevated plus maze test (*Nsf*^f/f^: n=10; *Nsf*^f/f^;D2R-Cre: n=10). (f) Startle response to a 120-dB acoustic stimulus (*Nsf*^f/f^: n=15; *Nsf*^f/f^;D2R-Cre: n=15). (g) Pre-pulse inhibition (PPI) performance (*Nsf*^f/f^: n=15; *Nsf*^f/f^;D2R-Cre: n=15). (h) Cumulative time spent interacting with a stranger mouse versus in an empty cage in the three-chamber social interaction test (*Nsf*^f/f^: n=13; *Nsf*^f/f^;D2R-Cre: n=11). Data are presented as mean±SEM. Student’s *t*-test; \*\**p*<0.01 (a, c, h), \**p*<0.05 (e).

However, time spent in the open arms did not differ significantly between the groups (*Nsf*^f/f^ mice: 81±37 s; *Nsf*^f/f^;D2R-Cre mice: 13±5 s; Student’s *t-*test, *p*=0.099; Fig. 5e). No significant differences were observed between the groups in startle response to a 120-dB acoustic stimulus (*Nsf*^f/f^ mice: 1 911±227 mV; *Nsf*^f/f^;D2R-Cre mice: 1 611±266 mV; Student’s *t-*test, *p*=0.42; Fig. 5f) or in PPI across various intensities (74 dB, *Nsf*^f/f^ mice: 34±3.25, *Nsf*^f/f^;D2R-Cre mice: 34±4.95, *p*=0.43; 80 dB, *Nsf*^f/f^ mice: 41±4.57, *Nsf*^f/f^;D2R-Cre mice: 43±4.8, *p*=0.32; 84 dB, *Nsf*^f/f^ mice: 46±6, *Nsf*^f/f^;D2R-Cre mice: 53±5, *p*=0.25; Student’s *t*-test) (Fig. 5g). In the three-chamber social interaction test, both groups spent significantly more time interacting with a cage containing a conspecific mouse (*Nsf*^f/f^ mice: 110±18 s; *Nsf*^f/f^;D2R-Cre mice: 75±16 s) than with an empty cage (*Nsf*^f/f^ mice: 23±5 s, *p*<0.001; *Nsf*^f/f^;D2R-Cre mice: 19±4 s, *p*=0.004; Student’s *t*-test) (Fig. 5h).

We further confirmed the hyperactivity and impulsivity of *Nsf*^f/f^;D2R-Cre mice at P28. *Nsf*^f/f^;D2R-Cre mice exhibited markedly increased locomotor activity in a novel environment, frequently running along the perimeter of the test box, unlike their littermate controls. The total travel distance was significantly greater in *Nsf*^f/f^;D2R-Cre mice (150 745±18 208 cm) than in controls (*Nsf*^f/f^ mice: 11 930±1 457 cm; *Nsf*^f/-^;D2R-Cre: 14 248±2 210 cm; *Nsf*^f/+^mice: 13 748±943 cm; Student’s *t*-test, *p*<0.001; Supplementary Fig. 3a, b). In the cliff avoidance test, *Nsf*^f/f^;D2R-Cre mice showed significantly shorter jump latency than did *Nsf*^f/f^ mice (*Nsf*^f/f^ mice: 410±10 s; *Nsf*^f/f^;D2R-Cre mice: 224±40 s; Student’s *t*-test, *p*<0.001; Supplementary Fig. 3c,d). During the 7-min session, 10% *Nsf*^f/f^;D2R-Cre mice and 0% *Nsf*^f/f^ mice jumped off within the first minute. By the end of the session, 90% *Nsf*^f/f^;D2R-Cre mice and only 7.6% *Nsf*^f/f^ mice had jumped off the platform (Supplementary Fig. 3e).

### Coadministration of MPH and the D2R agonist quinpirole (QNP) improved ADHD-like hyperactivity and impulsivity

*Nsf*^f/f^;D2R-Cre mice exhibited hyperactivity and impulsivity, hallmark features of ADHD. To explore potential treatments, we investigated whether MPH, a psychostimulant widely used in ADHD management, could ameliorate these behaviours in 4-week-old *Nsf*^f/f^;D2R-Cre mice. Locomotor activity was monitored in both *Nsf*^f/f^;D2R-Cre mice and control *Nsf*^f/f^ mice following MPH administration. MPH treatment alone did not significantly reduce hyperactivity in *Nsf*^f/f^;D2R-Cre mice (*Nsf*^f/f^;D2R-Cre mice: -30–0 min, 27 840.7±5 208.4 cm; 30–60 min, 25 747.8±6 437.2 cm; Student’s *t*-test, *p*=0.42; Fig. 6a,a’). Next, we assessed the combined effect of MPH and the D2R agonist QNP, aiming to compensate for the loss of striatal D2R-expressing cells. Following coadministration of MPH and QNP, a significant reduction in locomotion was observed in *Nsf*^f/f^;D2R-Cre mice (*Nsf*^f/f^;D2R-Cre mice: -30–0 min, 33 882.4±7 376.4 cm; 30–60 min, 5 242.1±5 016.4 cm; Student’s *t*-test, *p*=0.0004; Fig. 6b,b’). Treatment with QNP alone also produced a modest reduction in locomotion (*Nsf*^f/f^;D2R-Cre mice: -30–0 min, 17 529.9±2 761.5 cm; 30–60 min, 10 238.8±3 157.8 cm; Student’s *t-*test, *p*=0.049; Fig. 5c,c’). We further assessed impulsive behaviour in *Nsf*^f/f^;D2R-Cre mice following MPH and QNP treatment. Treated *Nsf*^f/f^;D2R-Cre mice displayed significantly longer jump latency than did their untreated counterparts (*Nsf*^f/f^;D2R-Cre mice: 224±40 s; treated *Nsf*^f/f^;D2R-Cre mice: 390±30 s; Student’s *t*-test, *p*=0.006). In contrast, *Nsf*^f/f^ mice exhibited the opposite response: treated *Nsf*^f/f^ mice exhibited a significantly shorter jump latency than did their untreated controls (*Nsf*^f/f^ mice: 410±10 s; treated *Nsf*^f/f^ mice: 174±59 s; Student’s *t-*test, *p*<0.001; Fig. 6d). During the 7-min session, the proportion of *Nsf*^f/f^;D2R-Cre mice that jumped off the platform within the first minute decreased from 10% (untreated) to 0% (treated). By the end of the session, the cumulative percentage of jumpers decreased from 90% to 11% following MPH and QNP co-administration (Fig. 6e). Conversely, *Nsf*^f/f^ mice displayed the opposite trend: by the end of the session, the percentage of jumpers increased from 7.6% in the untreated group to 70% after treatment. These findings suggest that co-administration of MPH and QNP may effectively alleviate hyperactivity and impulsive behaviour in *Nsf*^f/f^;D2R-Cre mice, offering mechanistic insights into potential therapeutic strategies for ADHD-like symptoms arising from D2R dysfunction.

**Fig. 6:**
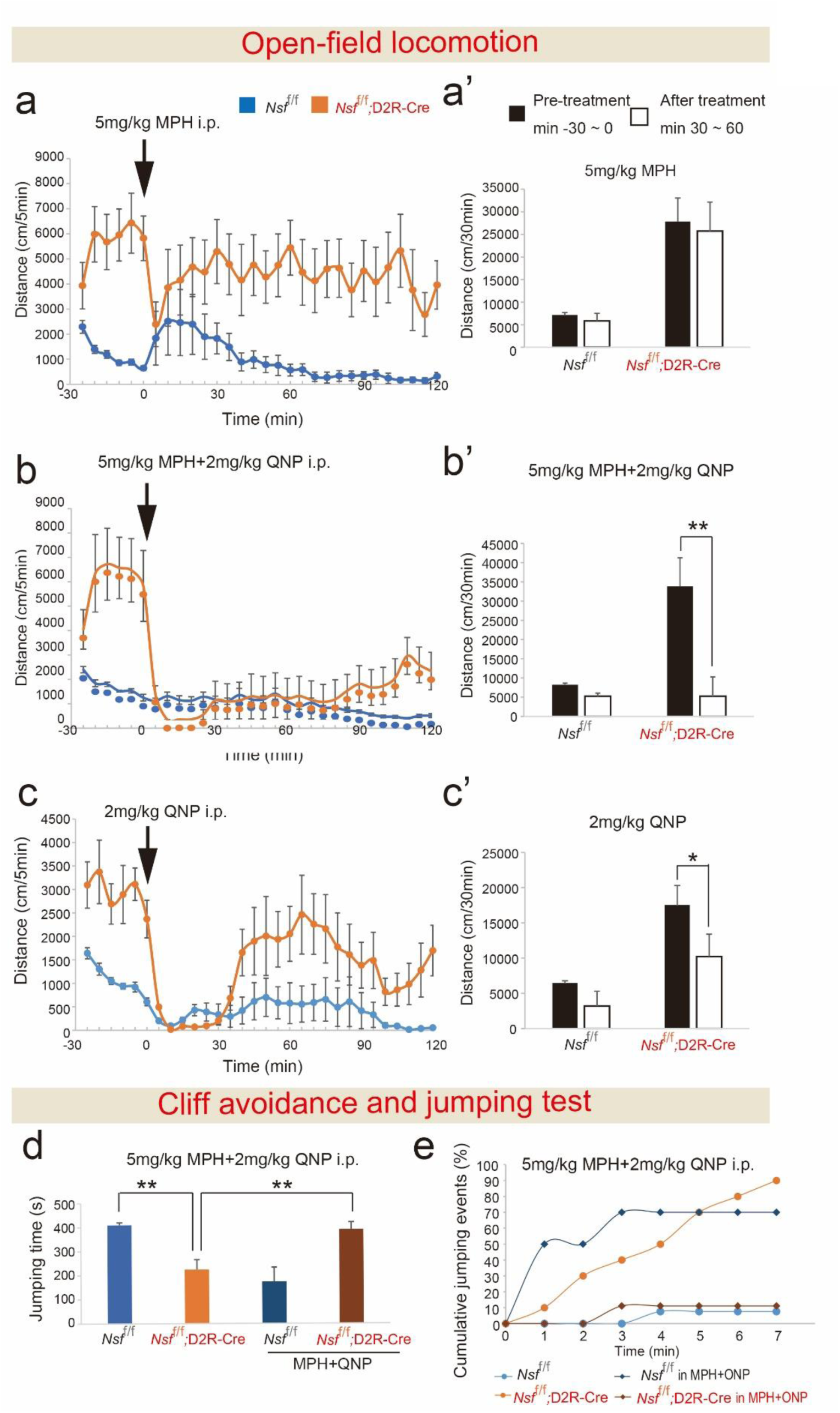
Coadministration of methylphenidate (MPH) and D2R agonist quinpirole (QNP) ameliorates locomotor hyperactivity and impulsivity in *Nsf*^f/f^;D2R-Cre mice. (a-c) Effects of MPH and QNP on locomotor activity in 4-week-old *Nsf*^f/f^;D2R-Cre mice exposed to a novel environment. Locomotor activity was monitored at 5-min intervals for 30 min before drug administration and again after intraperitoneal (i.p.) injection of either 5 mg/kg MPH (a), 5 mg/kg MPH combined with 2 mg/kg QNP (b), or 2mg/kg QNP alone (c). (a’-c’) Distance travelled was measured 30 min before and 30–60 min after drug administration. (a’) *Nsf*^f/f^ (n=8) and *Nsf*^f/f^;D2R-Cre (n=6) mice treated with MPH; (b’) both genotypes (n=8 per group) treated with MPH + QNP; (c’) both genotypes (n=9 per group) treated with QNP alone. (d,e) Effects of MPH + QNP on impulsivity in 4-week-old *Nsf*^f/f^ and *Nsf*^f/f^;D2R-Cre mice. Groups comprised untreated controls (*Nsf*^f/f^: n=13; *Nsf*^f/f^;D2R-Cre: n=11) and MPH+QNP-treated animals (*Nsf*^f/f^: n=10; *Nsf*^f/f^;D2R-Cre: n=9). (d) Latency to the first fall in the cliff avoidance reaction test. (e) Cumulative number of cliff-jumping events over 7 min. Data are presented as mean±SEM. Student’s *t*-test; \*\**p*<0.01 (a′, b′), \**p*<0.05 (c′).

## DISCUSSION

In this study, we generated conditional knockout mice lacking NSF in D2R-expressing cells. The *Nsf*^f/f^;D2R-Cre mice exhibited distinct behavioural and neurochemical abnormalities, including hyperactivity and impulsivity, loss of striatal D2R-expressing neurons, increased postnatal neuronal cell death, striatal shrinkage, and reduced dopamine concentrations in the striatum. Notably, their hyperactivity and impulsivity were ameliorated by the coadministration of MPH and QNP. Collectively, these findings suggest that NSF plays a neuroprotective role in D2R-expressing cells during postnatal development and that the loss of these cells contributes to ADHD-like behavioural phenotypes through impaired dopaminergic transmission in the striatum.

### Role of the NSF-D2R interaction in neuronal survival

Our study provides new insights into the essential role of the NSF-D2R interaction in neuronal survival during postnatal development. Previous studies indicate that NSF directly binds to the intracellular domains of D2R, regulating its intracellular trafficking, surface expression, and functional stability [10, 13]. This interaction is particularly important given D2R’s established function in modulating neuronal excitability and protecting neurons from excitotoxicity [12]. *In vitro* evidence has shown that D2R activation triggers anti-apoptotic signalling cascades, including the PI3K/Akt and ERK pathways, which are critical for mitigating glutamate-induced cytotoxicity [30]. These pathways are also involved in synaptic plasticity and learning, suggesting that NSF-D2R disruption could have lasting effects not only on survival but also on cognitive flexibility and behavioural regulation—core deficits observed in ADHD. To date, the physiological relevance of the NSF-D2R interaction *in vivo* has largely remained unexplored, although the neuroprotective role of D2R signalling has been highlighted in neurological diseases including Parkinson’s disease, ischemia, and epilepsy [31].

In *Nsf*^f/f^;D2R-Cre mice, we observed a significant increase in apoptosis, particularly in the striatum during postnatal development, accompanied by reduced striatal volume. This structural abnormality parallels neuroimaging findings in individuals with ADHD [17]. Our results demonstrate that NSF is indispensable for stabilising D2R and supporting its neuroprotective function *in vivo*. In the absence of NSF, D2R-expressing neurons exhibit heightened vulnerability to apoptosis, underscoring the role of NSF in maintaining D2R stability and anti-apoptotic signalling. These findings establish NSF as a key regulator of neuronal survival through its interaction with D2R. Consistent with this, D2R knockout models have confirmed the receptor’s role in preventing excitotoxicity-induced neurodegeneration [31, 32]. The concordance between D2R knockout and NSF-deficient phenotypes indicates that NSF acts upstream of D2R, to stabilise its neuroprotective capacity. Together, these results support a coherent mechanistic framework: NSF regulates D2R integrity; D2R activation prevents excitotoxicity; and thus, NSF loss indirectly leads to neurodegeneration. This mechanistic link aligns with psychiatric molecular models that attribute synaptic protein dysregulation to circuit-level dysfunction and behavioural pathology.

The striatum within the basal ganglia plays a crucial role in various processes, receiving dopaminergic input from the SN pars compacta (SNc) and VTA [33]. D2R, expressed in midbrain dopamine neurons, regulates dopamine synthesis, release, and reuptake [34–36]. We hypothesised that the neuroprotective effects of the NSF-D2R interaction extend to the midbrain. In *Nsf*^f/f^;D2R-Cre mice, dopamine levels were markedly decreased, likely due to loss of D2R-expressing neurons in the SNc. This observation emphasises the importance of the NSF-D2R interaction in sustaining dopamine production and secretion. The concurrent reductions in DAT and TH expression in *Nsf*^f/f^;D2R-Cre mice further indicate impaired dopaminergic function, as TH mediates dopamine synthesis and DAT enables dopamine reuptake. The combined decline in dopamine production, release, and reuptake highlights the broad impact of NSF-D2R loss on dopaminergic signalling.

### ADHD-like behaviours of *Nsf*^f/f^;D2R-Cre mice

The *Nsf*^f/f^;D2R-Cre mice exhibited ADHD-like behaviours, including hyperactivity and impulsivity. These traits were reflected in increased locomotor activity and impaired inhibitory control, corresponding to the core behavioural symptoms of ADHD. These findings indicate that the loss of D2R-expressing cells coincides with the emergence of ADHD-like behaviours, suggesting a potential link between D2R dysfunction and these behavioural abnormalities.

Dysregulation of D2R-mediated signalling has long been implicated in ADHD, supported by genetic and neuroimaging studies. For instance, the Taq A1 allele of the *D2R* gene has been associated with impulsive and compulsive behaviours in ADHD, likely due to reduced D2R availability in the basal ganglia [37, 38]. Similarly, positron emission tomography studies have reported diminished D2R/D3R density in key dopaminergic regions, including the nucleus accumbens and caudate nucleus, in individuals with ADHD [15]. Although the *D2R* gene has been proposed as a contributor to ADHD susceptibility, it is not as consistently implicated as *DRD4* or *DAT1*. The focus on other dopamine-related genes and the catecholaminergic system in general indicates that ADHD pathophysiology involves multiple genetic and neurochemical factors beyond D2R alone [39–42]. The *Nsf*^f/f^;D2R-Cre mouse model extends these findings by demonstrating that the loss of NSF-D2R-expressing cells can replicate these behavioural impairments, highlighting the translational relevance of this model for understanding dopaminergic contributions to ADHD.

Our research offers new perspectives on potential therapeutic strategies for ADHD. Although MPH remains a first-line treatment that primarily increases synaptic dopamine and noradrenaline levels, it does not directly target D2R dysfunction [43, 44]. Notably, approximately 30% of patients with ADHD show limited response to MPH, underscoring the necessity for alternative therapies [45]. Research indicates that the D2R subtype influences both hyperactivity and amphetamine response in ADHD models [46]. Drug-naïve children with ADHD exhibit elevated D2R availability, which normalises following MPH therapy [47]. Moreover, higher baseline D2R levels correlate with improved MPH response, particularly in reducing hyperactivity [47]. Genetic studies also link the TaqI A polymorphism of the *D2R* gene to ADHD, with the A1 allele and A1A1 genotype associated with ADHD in males [38. Collectively, these findings emphasise the pivotal role of D2R in ADHD pathogenesis and treatment efficacy, potentially opening avenues for innovative therapeutic strategies. Interestingly, combining MPH with a D2R agonist effectively mitigated hyperactivity in *Nsf*^f/f^;D2R-Cre mice, indicating a synergistic approach that tackles both dopamine availability and receptor-specific dysfunction.

The *Nsf*^f/f^;D2R-Cre mouse model serves as an important tool for investigating the underlying mechanisms of ADHD. The behavioural and neurochemical characteristics exhibited by these mice strongly support the crucial role of NSF-D2R interactions in the manifestation of ADHD-like symptoms. This innovative model provides researchers with a unique opportunity to delve into the pathophysiology of ADHD and potentially uncover new therapeutic approaches targeting D2R-associated dysfunctions.

### Limitations and future directions

Despite the insights gained, this study has several limitations. First, we could not directly confirm NSF deletion in D2R-expressing neurons due to early degeneration; future studies using inducible Cre systems or lineage tracing are needed. Second, our analyses focused on the striatum, even though D2R neurons exist in other regions. Although partial loss of TH-positive neurons was detected in the midbrain, the broader impact of NSF deletion on extra-striatal D2R populations remains unknown. Whole-brain mapping will clarify whether NSF function is universally required for D2R neuron maintenance. Third, the molecular mechanism linking NSF deficiency to apoptosis is not fully understood. Single-cell transcriptomic or proteomic analyses may reveal downstream pathways. Fourth, while D2R neuron loss, reduced dopamine, and ADHD-like behavior occurred in parallel, causality remains unproven. It is unclear whether restoring D2R function or dopamine tone rescues these behaviors. Finally, the mechanism behind the synergistic effect of methylphenidate and quinpirole remains unknown. Further investigation may clarify how these drugs compensate for NSF–D2R dysfunction. Addressing these limitations will help define NSF’s role in dopaminergic neuron survival and its broader relevance to ADHD pathophysiology.

In conclusion, our study underscores the critical role of NSF-D2R interactions in neuronal viability and demonstrates that their impairment results in neurophysiological and behavioural features resembling ADHD (Supplementary Fig. 4). Although ADHD is characterised by dopaminergic dysfunction involving multiple neurotransmitter receptors and transporters, including D2R, DAT, and dopamine itself, a comprehensive investigation of their combined influence on ADHD pathophysiology remains underexplored. Moreover, the notion that these neurotransmitter abnormalities might stem from a shared upstream cause has not been thoroughly examined.

To our knowledge, the *Nsf*^f/f^;D2R-Cre mouse model offers the first *in vivo* evidence demonstrating how the elimination of NSF-D2R-expressing cells contributes to decreased dopamine levels and emphasises the crucial role of D2R in maintaining striatal function. These insights enhance our comprehension of ADHD pathophysiology and highlight the potential for developing specific therapeutic approaches aimed at restoring dopaminergic equilibrium in the striatum.

## Acknowledgements

We are grateful to K, Iwata, N. Miyagoshi, Y. Sasaki, F. Yamamoto, R. Ogawa, S. Kanae, and I. Kumano for technical assistance and T. Taniguchi for secretarial assistance. We also thank NPO Biotechnology Research and Development for technical assistance. We are grateful to N. Heintz and C. Gerfen for donating D2R-Cre mice.

## Author Contributions

M-J. X. performed most experiments and wrote the manuscript jointly with K. Murata.

K. M. conducted *in situ* hybridisation experiments and co-wrote the manuscript with M-JX.

H. K. performed the AAV experiments.

Y. F. provided advice on *in situ* hybridisation.

N. U. supervised the project and co-wrote the manuscript with M-JX.

H. M. conceived and directed the project and co-wrote the manuscript with M-JX.

All authors provided critical feedback and contributed to the final version of the manuscript.

## Availability of Data and Materials

All data generated or analysed during this study are included in this published article and its supplementary information files. Additional datasets are available from the corresponding author upon reasonable request.

## Supplementary Information

Supplementary information is available at MP’s website

## SUPPLEMENTARY INFORMATION

### SUPPLEMENTARY METHODS

#### Genotyping polymerase chain reaction (PCR)

Genomic DNA was extracted from ear punches, and genotyping was performed using PCR. *For Nsf^f/+^*mice, the following primers: 5′-AGGAGCCTTCTGGATGACCT and 5′-GGATCCATGGCTTCAATGTC, yielding an 853-bp product corresponding to the transgenic/floxed allele. For D2R-cre mice, the primers: 5′-GTGCGTCAGCATTTGGAGCAA and 5′-CGGCAAACGGACAGAAGCATT, producing a 700-bp product indicative of the transgenic *D2R-Cre* allele. The PCR product was sequence-verified for accuracy.

#### Stereotaxic surgery and adeno-associated virus (AAV) injection in the striatum

Mice were anaesthetised by intraperitoneal injection of a mixture containing medetomidine (0.75 mg/kg), midazolam (4 mg/kg), and butorphanol (5 mg/kg), and secured in a stereotaxic frame (Narishige, Tokyo, Japan). To prevent corneal drying, an ophthalmic ointment (Tarivid, Santen, Osaka, Japan) was applied to the eyes. A small cranial window was made above the left striatum (0.5 mm anterior to bregma, 1.75 mm lateral to the midline). A mixture of viral vectors, AAV5-human synapsin (hSyn)-EGFP (#50465, titre: 8.4 × 10^12^ vg/mL) and AAV5-hSyn-DIO-mCherry (#50459, titre: 8.4 × 10^12^ vg/mL) (Addgene, Watertown, USA) was combined at a 1:1 ratio and injected into the striatum (depth: 2.0 mm from the dura; volume: 300 nL; infusion rate: 100 nL/min) using a 10 μL Hamilton syringe connected to an infusion pump (UMP-3, World Precision Instruments, FL, USA). After infusion, the needle was left in place for 10 min before being slowly withdrawn, and the scalp was sutured. At the end of surgery, mice received an intraperitoneal injection of atipamezole (0.75 mg/kg) to facilitate recovery from anaesthesia. Four weeks after viral injection, mice were deeply anaesthetised with isoflurane and perfused transcardially with phosphate-buffered saline, followed by 4% paraformaldehyde. Brains were removed, post-fixed in 4% paraformaldehyde, and sectioned coronally at 50 μm using a vibratome (Dosaka EM Co., Ltd., Kyoto, Japan). Sections were mounted using DAPI-containing medium and imaged with a BZ-X800 All-in-One Fluorescence Microscope (Keyence, Tokyo, Japan).

### Behavioural tests

#### Open-field locomotion

The open-field test was conducted to assess spontaneous locomotor activity and exploratory behaviour in a novel environment, as previously described [1]. Adult mice were individually placed in a 48 × 48 cm² open-field arena (MELQUEST Co., Toyama, Japan). At the beginning of the test, each mouse was positioned in the front right corner of the arena and allowed to explore freely for 120 min. Their movements were continuously recorded and analysed, as reported previously [1]. For drug treatment experiments, mice were placed in the same arena and monitored at 5-min intervals for 30 min to assess baseline activity. Following this period, 4-week-old mice received intraperitoneal (i.p.) injections of one of the following treatments: 5 mg/kg methylphenidate (MPH), 2 mg/kg quinpirole (QNP), or a combination of 5 mg/kg MPH and 2 mg/kg QNP. Immediately after injection, mice were returned to the arena, and behaviour was recorded for another 90 min.

#### Cliff avoidance and jumping test

The cliff avoidance and jumping test was conducted to assess impulsivity-like or risk-taking behaviour in mice, as previously described [2, 3]. Offspring aged over 6 months were used for the experiment. The test platform consisted of an inverted glass beaker with a diameter of 13.5 cm and a height of 20.3 cm, ensuring that its height exceeded twice the average body length of the mice to simulate a perceived ‘cliff’. Each mouse was gently placed at the centre of the platform to minimise initial stress. Behaviour was observed for up to 7 min or until the mouse jumped off the platform, whichever occurred first. The latency to jump (the time from initial placement to the first jump) was recorded for each mouse. In addition, the cumulative number of mice that had jumped off the platform was recorded at the end of each minute. Mice that did not jump within the 7-min session were included in the total sample size for percentage calculations. At each time point, the percentage of cumulative jumping events was calculated using the following formula: percentage of cumulative jumping events = (T/TN) ×100 Where N is the number of mice that had jumped off by the specific minute, and TN is the total number of mice tested in the session [2, 3]. For drug treatment experiments, 4-week-old mice received i.p. injections of a combination of 5 mg/kg MPH and 2 mg/kg QNP. After a 30-min interval, the mice underwent the cliff avoidance and jumping test using the same protocol as described above.

#### Elevated plus maze (EPM) test

The EPM test was conducted, as previously described [4], to evaluate anxiety-like behaviour in mice. The EPM apparatus consisted of two perpendicular runways forming a plus (‘+’) shape, with two open arms (without walls) and two closed arms (with walls). Each arm measured 25 cm in length and 5 cm in width, and the closed arms had walls 25 cm high (Lab&design, Tokyo). The maze was elevated 50 cm above the floor level to induce a natural conflict between exploration and the fear of open spaces. At the beginning of each trial, each mouse was placed in the central square (intersection of the arms), facing one of the closed arms, and allowed to freely explore the maze for 5 min. The time spent in the open arms was recorded as a primary measure of anxiety-like behaviour. Behavioural data were automatically recorded and analysed using a video tracking system (Smart 3.0, Panlab, Barcelona, Spain).

#### Prepulse inhibition (PPI) test

PPI of the auditory startle reflex is a widely recognised operational measure of sensorimotor gating. PPI of the acoustic startle response was assessed using a startle response system (SR-LAB, San Diego Instruments, CA, USA). All experiments were conducted in a sound-attenuated, ventilated chamber, with events recorded and controlled using the MED Associates software (Startle Reflex package). Adult mice were individually placed in cylindrical Plexiglass holders mounted on a platform equipped with a piezoelectric accelerometer to detect startle-induced movements. After a 5-min acclimation period in the presence of background white noise (69 dB), mice were presented with a series of trials comprising: pulse-alone trials: a single 120-dB startle stimulus (40 ms); prepulse + pulse trials: a 20-ms prepulse sound at 74, 80, or 84 dB followed by the 120-dB pulse after a 100-ms interval; prepulse-alone trials: prepulse only without the pulse; and no-stimulus trials: background noise only. Each session included 60 trials. The magnitude of the startle response was recorded for each trial. The degree of PPI was calculated using the following formula: The percentage PPI induced by each prepulse intensity was calculated as [1-(startle amplitude on prepulse trial)/(startle amplitude on pulse alone)]×100%. Each animal’s response was averaged across trials at each prepulse intensity. Higher PPI values indicated greater sensorimotor gating ability.

#### Three-chamber test

The three-chamber test was performed as previously described [1]. The apparatus consisted of a rectangular box divided into three compartments and was covered with a lid equipped with an infrared video camera (TimeCSI2; Ohara & Co., Tokyo, Japan). In Session I (habituation), the subject mouse was placed in the central chamber of the empty apparatus and allowed to freely explore all three chambers for 10 min. In Session II (sociability test), an unfamiliar C57BL/6N female mouse (the ‘stranger mouse’) was placed in a cage within one of the side chambers. The subject mouse was then allowed to freely explore the apparatus for another 10 min. The time the subject mouse spent in direct contact with the cage containing the stranger mouse was recorded as an index of sociability.

---

**Supplementary Fig. 1:**
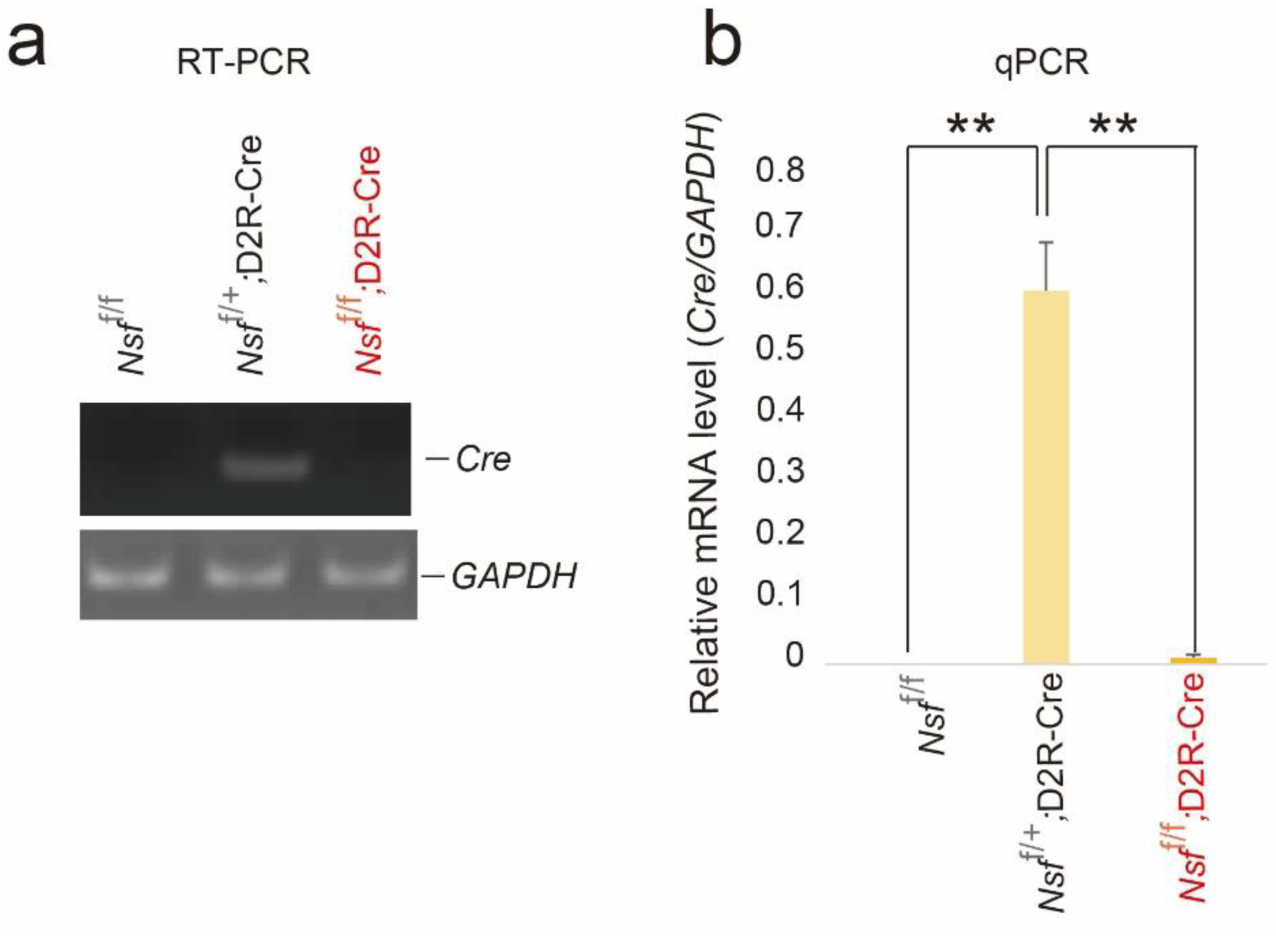
Reduction in *Cre* expression in the striatum of *Nsf*^f/f^;D2Rcre mice. (a) *Cre* mRNA expression in the striatum of *Nsf*^f/f^, *Nsf*^f/+^;D2R-Cre mice, and *Nsf*^f/f^;D2R-Cre mice, as detected using RT-PCR. (b) *Cre* mRNA expression was quantified via qPCR, with *Gapdh* as an internal reference (n=7). Data are presented as mean ± SEM. Student’s *t*-test; \*\**p*<0.01.

**Supplementary Fig. 2:**
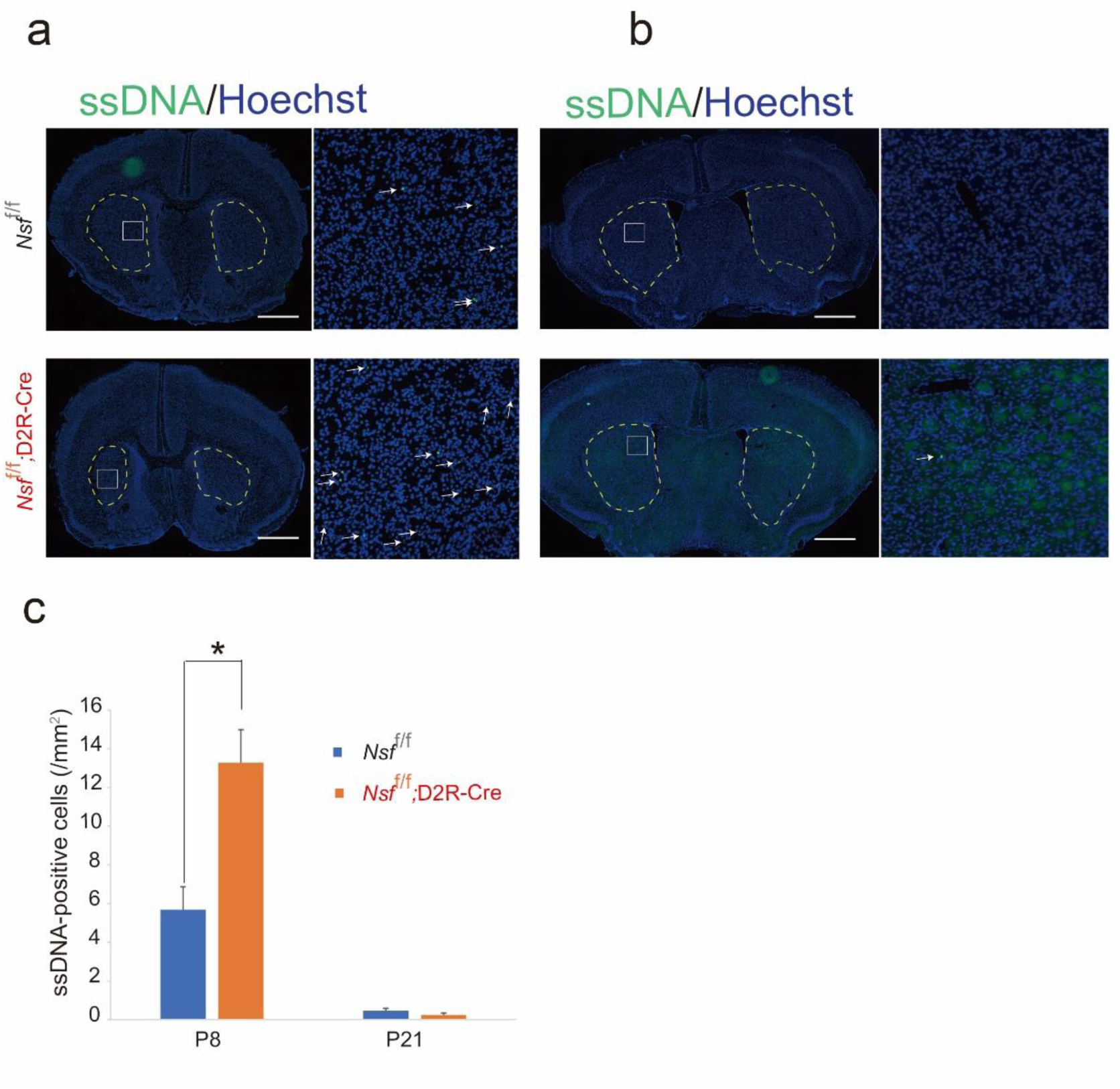
Increased cell death during postnatal development in *Nsf*^f/f^;D2R-Cre mice. ssDNA expression was detected via immunofluorescence, with all cells visualised using Hoechst staining (blue). Scale bar=1 mm. (a) Postnatal day 8 (P8) striatal sections from *Nsf*^f/f^ (n=12) and *Nsf*^f/f^;D2R-Cre (n=10) mice. (b) P21 striatal sections from *Nsf*^f/f^(n=20) and *Nsf*^f/f^;D2R-Cre (n=10) mice. (c) Quantification of ssDNA-positive cells per mm^2^ in the striatum of *Nsf*^f/f^ and *Nsf*^f/f^;D2R-Cre mice. Data are presented as mean±SEM. Student’s *t*-test; \*\**p*<0.01.

**Supplementary Fig. 3:**
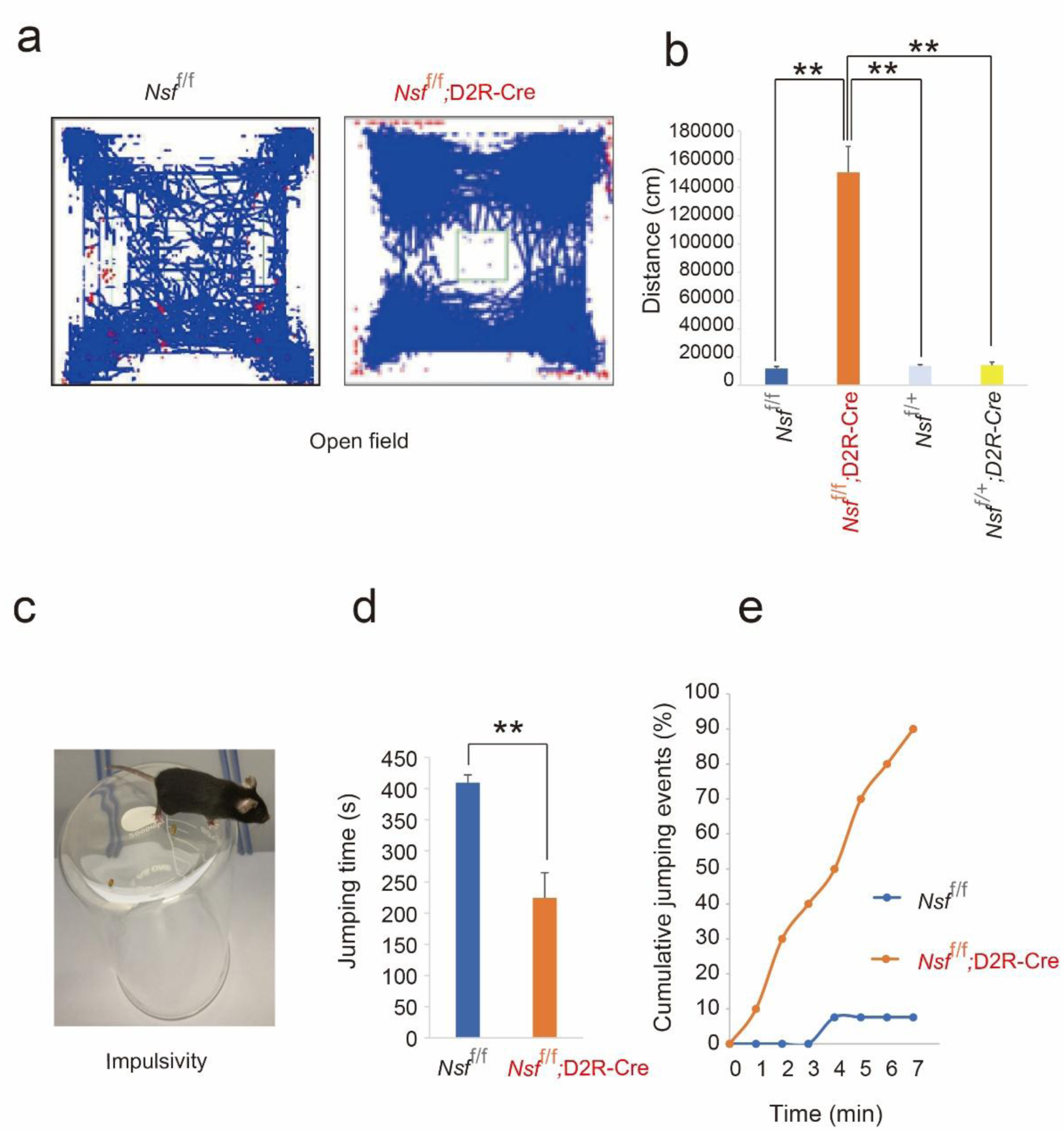
Hyperactivity and impulsivity in *Nsf*^f/f^;D2R-Cre mice. (a) Representative movement traces showing the locations of *Nsf*^f/f^ and *Nsf*^f/f^;D2R-Cre mice during the open field test. (b) Total distance travelled by *Nsf*^f/f^;D2R-Cre mice at 4 weeks of age (*Nsf*^f/f^, n=18; *Nsf*^f/f^;D2R-Cre, n=16; *Nsf*^f/+^, n=8; *Nsf*^f/+^;D2R-Cre, n=7). (c) Schematic diagram of the cliff avoidance reaction test paradigm. Mice were gently placed on a raised platform. (d) The time from initial placement on the platform until the first fall was measured for each group (*Nsf*^f/f^, n=13; *Nsf*^f/f^;D2R-Cre, n=10). (e) Cumulative frequency of cliff-jumping events over a 7-min period for *Nsf*^f/f^ (n=13) and *Nsf*^f/f^;D2R-Cre (n=10) mice. Data are presented as mean±SEM. Student’s *t*-test; \*\**p*<0.01 (b, d).

**Supplementary Fig. 4:**
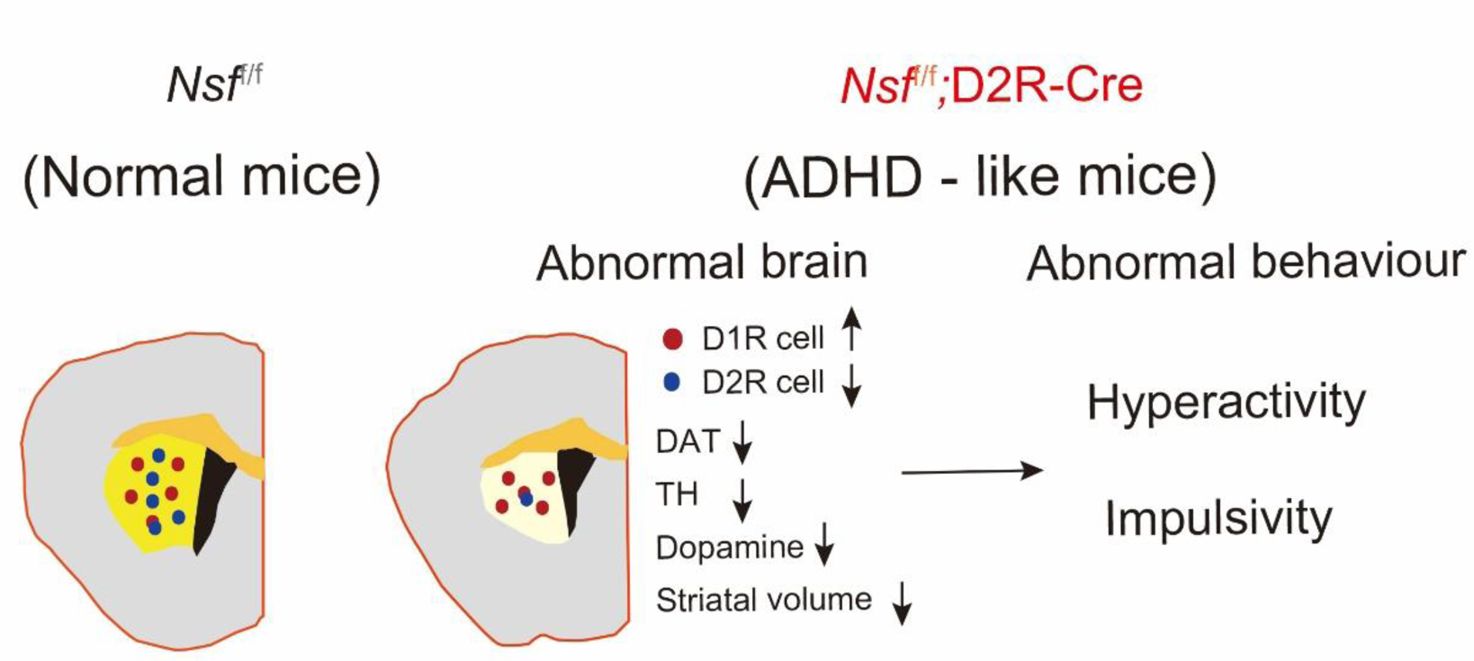
Graphic summary. Schematic illustration of the potential mechanisms by which *Nsf* deficiency induces neurophysiological and behavioural abnormalities resembling ADHD in mice, mediated by the loss of D2R-expressing cells and abnormal amino acid metabolism.

